# ppGpp is a bacterial cell size regulator

**DOI:** 10.1101/2020.06.16.154187

**Authors:** Ferhat Büke, Jacopo Grilli, Marco Cosentino Lagomarsino, Gregory Bokinsky, Sander Tans

## Abstract

Growth and division are central to cell size. Bacteria achieve size homeostasis by dividing when growth has added a constant size since birth, termed the “adder” principle, by unknown mechanisms [1–4]. Growth is well known to be regulated by ppGpp, which controls anything from ribosome production to metabolic enzyme activity and replication initiation, and whose absence or excess can induce the stress response, filamentation, and yield growth-arrested miniature cells [5–8]. These observations raise unresolved questions about the relation between ppGpp and size homeostasis mechanisms during normal exponential growth. Here, to untangle effects of ppGpp and nutrients, we gained control of cellular ppGpp by inducing the synthesis and hydrolysis enzymes RelA and Mesh1. We found that ppGpp not only exerts control over the growth rate, but also over cell division and hence the steady state cell size. The added size responds rapidly to changes in the ppGpp level, aided by transiently accelerated or delayed divisions, and establishes its new constant value while the growth rate still adjusts. Moreover, the magnitude of the added size and resulting steady-state birth size correlate consistently with the ppGpp level, rather than with the growth rate, which results in cells of different size that grow equally fast. Our findings suggest that ppGpp serves as a critical regulator that coordinates cell size and growth control.

## Results

### Control of ppGpp synthesis and hydrolysis

To study the relation between ppGpp and cell growth and division, two enzymes were used: the catalytic domain of the *E. coli* ppGpp synthesis enzyme RelA (RelA*) [9,10], and the ppGpp hydrolysis enzyme Mesh1 from *Drosophila melanogaster* [11,12], which were fused to YFP and CFP respectively. The former was inducible by doxycycline (Dox) and the latter by isopropyl-β-D-thiogalactopyranoside (IPTG) (Fig. 1A). As a control, we showed that the expression of a fluorescent protein alone did not significantly affect cell size or growth rate (Supp. Fig. 1E-F). We characterized this co-expression system in a ppGpp^0^ strain (*ΔrelA, ΔspoT*) that cannot produce ppGpp. In minimal medium lacking amino acids, growth was undetectably low in absence of RelA* induction, consistent with previous reports [8,12]. ppGpp is then required to activate amino acid biosynthesis operons [13,14]. However, growth became exponential if both RelA* and Mesh1 we co-expressed (Supp. Fig. 2A). These findings confirm that balanced synthesis and hydrolysis can achieve the constant ppGpp levels that are critical to normal exponential growth. If RelA* and Mesh1 indeed counteract in ppGpp production, then the same growth rates should be achievable by increasing both in parallel, as the additional synthesis by RelA* can then be canceled by the additional hydrolysis by Mesh1. The data indeed showed similar growth profiles for different combinations of RelA* and Mesh1 expression; with both either at lower levels, or both at higher levels (Supp. Fig. 2A).

**Figure 1.**
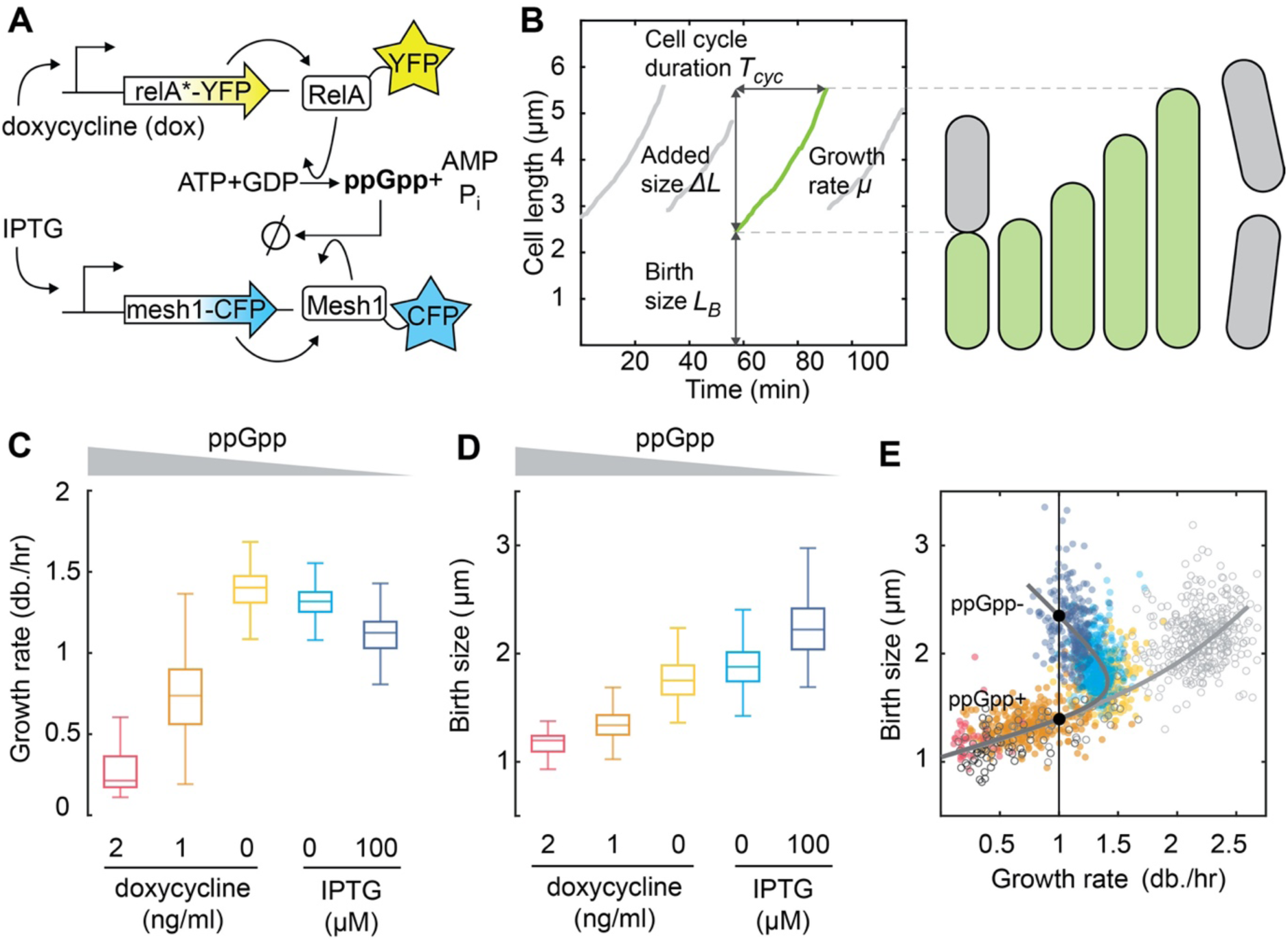
ppGpp exerts cell size control. **(A)** Scheme to control the cellular ppGpp concentration. A RelA truncate from *E. coli* (*relA\**), which synthesizes ppGpp, is fused to YFP and induced by Dox. The ppGpp hydrolysis enzyme Mesh1 is fused to CFP and induced by IPTG. See Fig. S1 and S2 for characterization of these constructs. **(B)** Measured length of single cells grown in a microfluidic device. For each cell cycle we quantify: size (length) at birth (*L*_*B*_), cell cycle duration (*T*_*cyc*_), added size (*ΔL*), and the growth rate (*μ*) by exponential fitting. **(C)** Growth rate for decreasing ppGpp levels. ppGpp increases above basal levels when inducing *relA\** with Dox and decreases below when inducing *mesh1-CFP* by IPTG, in *WT* (*relA+, spot+*) cells. Left to right: *N* = 42, 316, 257, 403, 241 cell cycles. *μ* peaks at basal (endogenous) ppGpp levels, and then decreases. **(D)** Birth size for decreasing ppGpp levels. Conditions as in panel C. *L*_*B*_ increases continuously while *μ* decreases for below-basal ppGpp levels. The *L*_*B*_ thus follows the ppGpp trend rather than that of *μ*. **(E)** Birth length against growth rate. Closed circles: single cell cycles in minimal media, colors and conditions as panels C and D. Drawn lines are guides to the eye. For below-basal ppGpp, slower growing cells are larger, owing to an inversion of the growth law. Cells of different size can thus have the same growth rate (black dots). Open circles: single cell cycles in rich media, with 2 ng/ml, 1 ng/ml, or 0 ng/ml Dox, and N = 66, 35, 336 cell cycles (dark to light gray).

### ppGpp exerts cell size control

We studied the effects of ppGpp at the single-cell level using a microfluidic chip that allowed media exchange, phase contrast and fluorescence microscopy, and cell-tracking algorithms [15,16]. We determined the length at birth (*L*_*B*_) and division (*L*_*D*_), the cycle duration (*T*_*cyc*_) and exponential growth or elongation rate (*µ*) for each cell cycle, and RelA* and Mesh1 enzyme concentrations, as quantified by the mean fluorescence per pixel (Fig. 1B, Supp. Fig. 1A-D). Here, we expressed either RelA*-YFP or Mesh1-CFP at moderate levels in the *WT* background (*relA*+ and *spoT+*), in order to produce minor deviations in the ppGpp concentration, from above to below basal levels, while maintaining balanced exponential growth (Fig. 1C, D).

The (population-mean) trend in the growth rate *µ* showed an optimum while the birth size *L*_*B*_ went up monotonically, as ppGpp decreased from above to below basal levels (Fig. 1C, D). The relation between *L*_*B*_ and *µ* (Fig. E) contained a number of intriguing features. First, as ppGpp decreased, both *L*_*B*_ and *µ* increased initially (Fig. 1E), in agreement with the well-known finding that faster growing cells are larger. However, decreasing ppGpp further led to an inverted trend, in which slower growing cells are larger (Fig. 1E). This deviation began at near-endogenous ppGpp levels (Fig. C-E). A counter-intuitive consequence of this inversion is that excursions above and below this endogenous ppGpp level lead to cells that differ in size but grow equally fast (Fig. 1E, two black dots on vertical line). The same trends were observed for the ppGpp^0^ strain with RelA* and Mesh1 (Supplementary Fig 2B-C).

The data are thus inconsistent with models in which ppGpp is a regulator of growth, and in turn, growth sets cell size, as described by the general growth law [1,17–19]. In such *hierarchical* models, *L*_*B*_ would follow the increase-optimum-decrease trend observed for *µ* (Fig. 1C). Instead, the monotonic increase of *L*_*B*_ with increasing ppGpp (Fig. 1D) suggested that ppGpp affects *L*_*B*_ in a way that is not mediated by *µ*. If ppGpp indeed modulates the size of cells in a *µ*-independent manner, we surmised it should play a role in the adder mechanism, which maintains the cell size constant against stochastic variations in birth size.

### ppGpp dynamically controls added cell size

In order to investigate the effect of ppGpp on the added size, we quantified the added length (*ΔL*) each cell cycle. First, we found that the adder principle was obeyed at all ppGpp concentrations: for the different levels of Dox and IPTG induction, *ΔL* was birth-size independent (Fig. 2A). In line with previous adder principle observations, we find that *T*_*cyc*_ rather than *µ* is modulated to achieve a constant *ΔL*, as larger-born cells divide sooner (Fig. 2B). Indeed, *ΔL* increased monotonically with decreasing ppGpp (Supplementary Fig. 2D), and thus did not follow the trend observed for *µ* (Fig. 1C). These data indicated that the added size correlated with population-average ppGpp concentration rather than the rate of growth.

**Figure 2.**
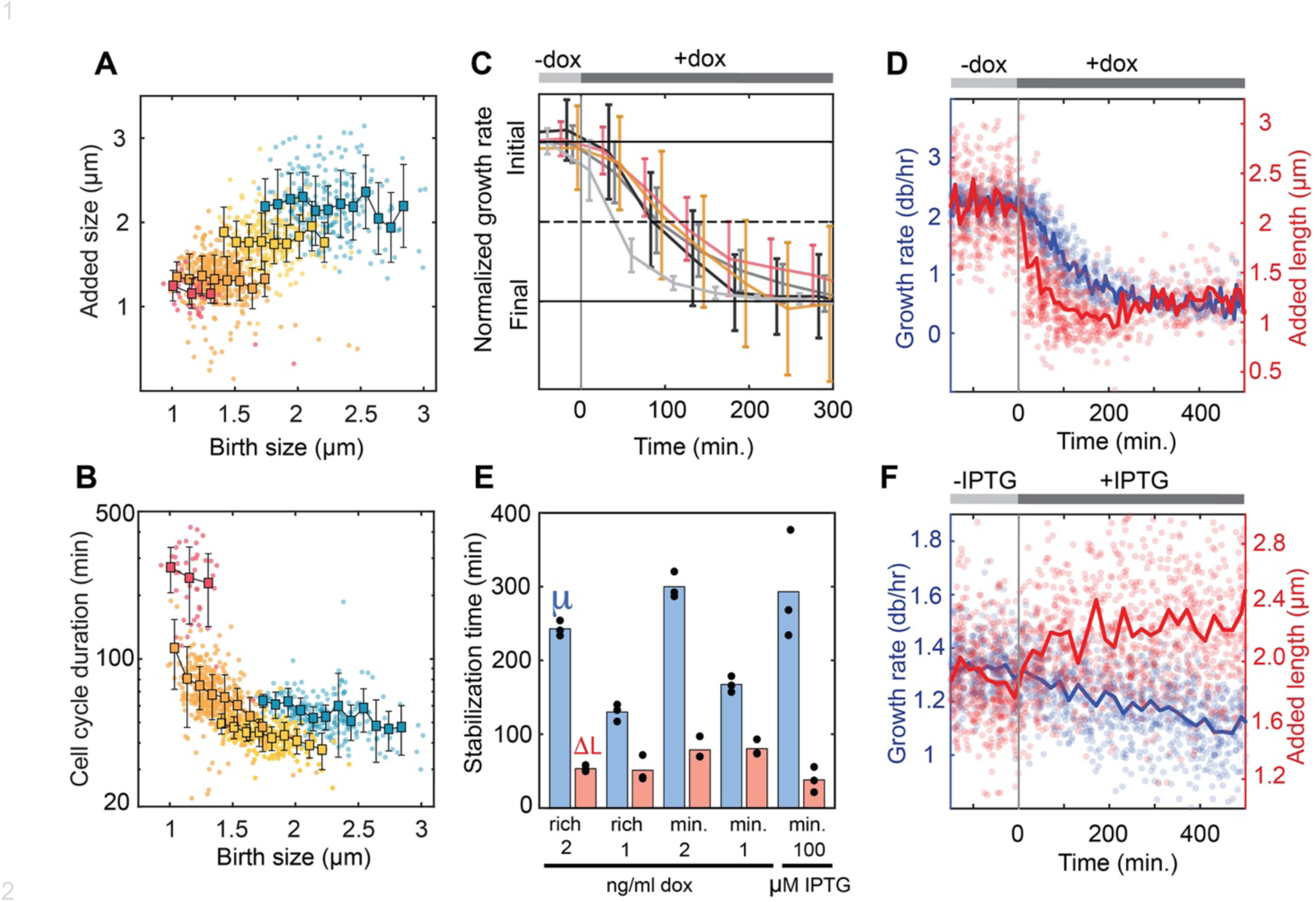
ppGpp dynamically controls added cell size. **(A)** Cell length added (*ΔL*) per cell cycle against birth length (*L*_*B*_) of that cycle, for different constant ppGpp levels. Dots are single cell cycles, squares are means for *L*_*B*_ bin, bars are s.e. Left to right: clouds for decreasing ppGpp levels of ppGpp, starting with Dox in ng/ml: 2 (red), 1 (orange), 0 (yellow), and then 100 uM IPTG (blue). *N* = 42, 316, 257, 241 cell cycles. For each cloud, *ΔL* is constant for different *L*_*B*_, consistent with “adder” principle. **(B)** Cell cycle duration (*T*_*cyc*_) against birth length (*L*_*B*_) of that cycle. When *L*_*B*_ is smaller, *T*_*cyc*_ is larger, indicating it is modulated as cells compensate for stochastic variations in *L*_*B*_, which is consistent with the adder principle. Colors and conditions as in panel A. **(C – F)** Cells during ppGpp shift, from basal to above and below basal levels. **(C)** Growth rate (*μ*) during ppGpp increases, by RelA* induction with Dox, in rich and minimal media. *μ* response timescale for these various conditions is assessed by normalizing rate to initial (pre-shift) and final (post-shift) value. Top bar indicates Dox induction. In minimal media, shift from 0 ng/ml Dox to: 2 (pink), and 1 (orange). In rich media, shift from 0 ng/ml to: 1 (dark gray), 2 (intermediate gray), and 10 (light gray). Bars are s.e.m., averaged over multiple cell cycles. Curves show similar adaptation time for moderate shifts, in contrast to the large 10 ng/ml shift, which is significantly faster, and serves as a control: growth indeed arrests at high ppGpp levels, as studied before, in line with the stringent response. **(D)** Growth (*μ*) and added size (*ΔL*) during ppGpp increase. Circles are single cell cycles, lines are moving averages, for shift from 0 to 2 ng/ml Dox, in rich medium. *ΔL* responds faster than *μ* and stabilizes to post-shift value while *μ* still varies. **(E)** Stabilization time for *μ* (blue) and *ΔL*(red). See methods for quantification approach. Each black dot shows the results of the analysis from 1/3^rd^ of the available data points. *ΔL* responds faster than *μ* under all moderate shifts. **(F)** Growth (*μ*) and added size (*ΔL*) during ppGpp decrease. Circles are single cell cycles, lines are moving averages, for shift from 0 to 100 μM IPTG Dox, in minimal medium.

Next, we considered how shifts in ppGpp concentration affect cell size and its control dynamically. Within the microfluidic flow-cell, we followed individual cells as they were exposed to a shift from basal ppGpp concentrations (-Dox) to different levels of RelA* induction (+Dox), in various growth media. First, the growth response underscored the important differences between the moderate induction that we focus on here (1 and 2 ng/ml) and the strong induction that mimics the stringent response (10 ng/ml Dox), with strong induction yielding a rapid growth arrest while moderate induction led to an approximately two-fold slower decrease in growth rate and allowed exponential growth to continue (Fig. 2C).

Notably however, the added size did respond rapidly even to low RelA* induction levels. Added size *ΔL* decreased halfway at about 25 min., and subsequently reached its final value at about 55 min (Fig. 2D, red trace). In contrast, *µ* responded substantially slower, and required about 100 min. to decrease halfway and over 300 min. to stabilize (Fig. 2C, blue trace). A similar pattern of rapid *ΔL* and slow *µ* responses were observed for different media and Dox induction levels (Fig. 2E), as well as for Mesh1-CFP induction (Fig. 2F). Consistently, *ΔL* increased more rapidly than *µ* decreased upon Mesh1-CFP induction (Fig. 2F).

The data show a temporal order in which *ΔL* responds to ppGpp deviations prior to *µ*. Indeed, *ΔL* typically has already stabilized to its post-shift level when *µ* has decreased only half-way from pre-induction to post-induction level (Fig. 2D-F). These data support the idea that *µ* and *ΔL* are decoupled, and further extend the notion that ppGpp affects cell size independently from its role in growth rate control. We hypothesize that a change in ppGpp concentration sets a new added size by affecting the division rate, which in turn results in a new steady-state birth size.

### Division accelerates transiently to achieve constant added size

We used mathematical modeling to understand the interplay between growth, division, and size, and compare different possible scenarios. Current size homeostasis models consider constant growth conditions, as well as a strict coupling between *μ* and *ΔL* [2,3,20,21]. We first tested a hierarchical model (Fig. 3A), which thus preserves the *μ*-*ΔL* coupling as growth conditions change. This model takes the observed initial and final values of *μ* and *ΔL* as input, lets *μ* decrease exponentially with the observed rate, and averages over multiple resulting simulated stochastic trajectories in *ΔL* and division rate 1/*T*_*cyc*_. The results of the hierarchical model (Fig. 3A) appeared inconsistent with the experimental observations (Fig. 3B). In particular, *ΔL* and *µ* decrease at similar rates in this model, while *L*_*B*_ decreases slower than *µ* (Fig. 3A), while the experiments show that both *ΔL* and *L*_*B*_ decrease faster than *µ* (Fig. 3B). Notably, the experimental data also indicate a transient increase in 1/*T*_*cyc*_ before it decreases (Fig. 3B), unlike the hierarchical control model (Fig. 3A).

**Figure 3.**
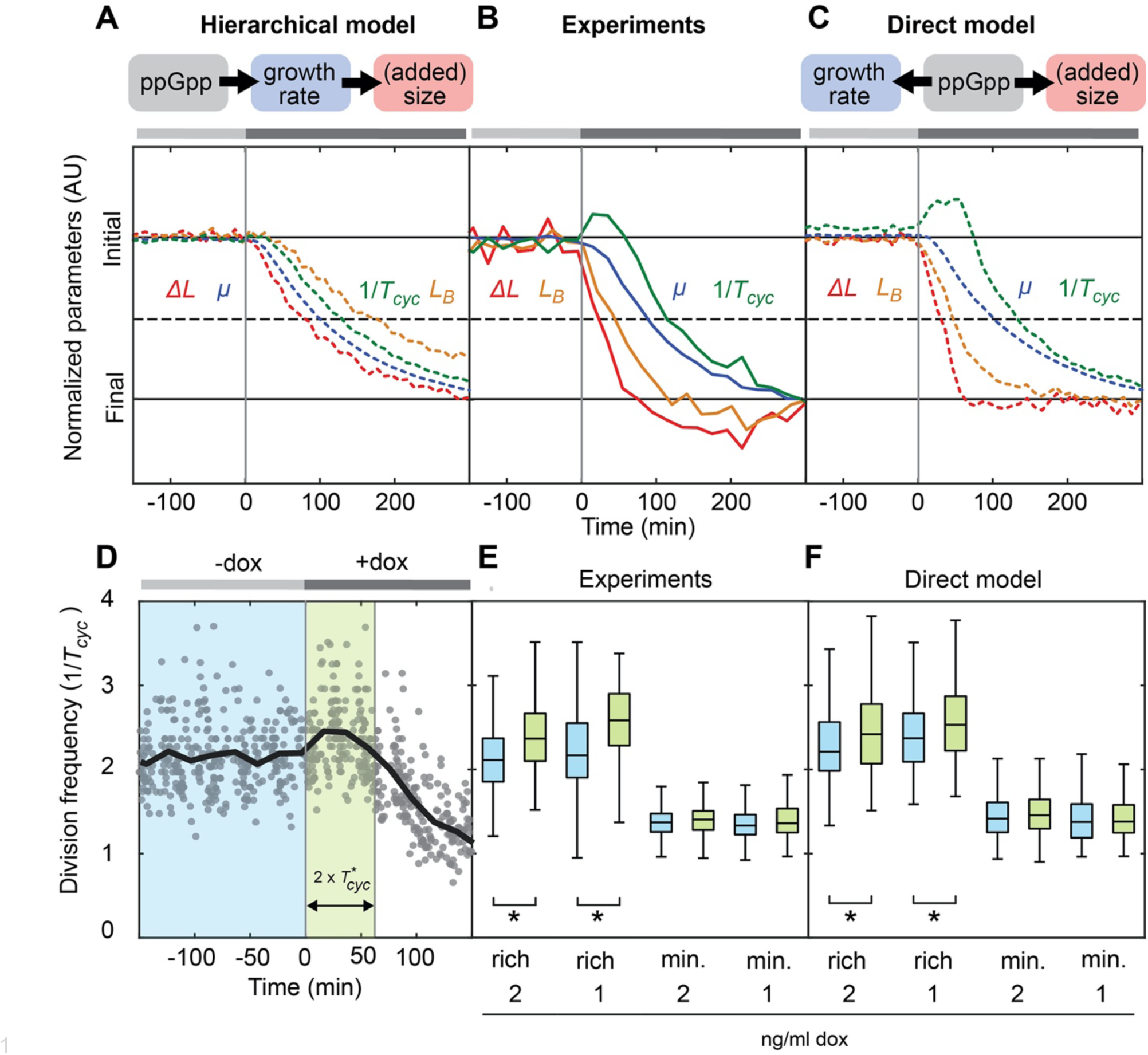
Division accelerates transiently to achieve constant added size. **(A)** Predictions of hierarchical model, in which ppGpp affects the growth rate (*μ*), and size in turn adjusts to the growth rate. To compare response speeds, indicated quantities are normalized to initial (pre-shift) and final (post-shift) values. Top bar indicates ppGpp increase. **(B)** Experimental data. Conditions and display as in panel A. **(C)** Predictions of direct model, in which ppGpp exerts control over division and hence (added) size independently of *μ*. Conditions and display as in panel A. Experiments agree with direct model in terms of the temporal order of the responses and the transient acceleration of divisions. **(D)** Quantification of transient effects in the division frequency (1/*T*_*cyc*_). Data is averaged in purple and green zones to obtain data in panels E and F. Circles are single cell cycles for shift from 0 to 2 ng/ml Dox, in minimal medium. Drawn line is moving average. **(E)** Pre- and post-shift division frequency (1/*T*_*cyc*_), as defined in panel D. In rich media, a ∼15% increase is observed between the blue and green zones. Star: *p* < 0.01. (N = 336, 186, 132, 64, 257, 191, 355, 214 left to right) **(F)** Direct model reproduces transient acceleration in rich media but not in minimal media, as seen in the experiments (N = 300, randomly taken from the simulations before induction, and N = 150 after induction).

Next, we considered a model of *direct* ppGpp control (Fig. 3C), which allows *μ* and *ΔL* to respond to ppGpp changes via independent routes and timescales. In this model, as also observed in the data (Fig. 2D), the mean *ΔL* responds directly by decreasing linearly to its post-shift value in about two pre-shift cell-cycles (note that division events in the averaged lineages are not synchronized), while *μ* decreases in the same way as in the previous model (Supp. Fig. 3A). The direct control model reproduces many features of the experimental data. Specifically, *L*_*B*_ responds slower than *ΔL* but faster than *µ*, and surprisingly, 1/*T*_*cyc*_ showed a transient increase (Fig. 3B and C).

This increase in 1/*T*_*cyc*_ may appear paradoxical, as it must decrease ultimately to match the lower *μ*. However, as illustrated by the model, it is a logical consequence of the *μ* and *ΔL* speed differences: *ΔL* reaches the lower post-shift value rapidly, while *μ* is still close to its high pre-shift value. With a comparatively high *μ*, the post-shift *ΔL* is achieved early, and hence divisions are early as well. Yet, it is indeed notable that the division rate appears readily modulated upwards and downwards. Together with the observation that *ΔL* is already stable at its new value while *µ* and 1/*T*_*cyc*_ are still varying towards theirs (Fig. 3B), these findings support a picture in which the cells act to realize their target *ΔL* (set in part by the concentration of ppGpp), and consequently *L*_*B*_, regardless of the rate of growth, and thus yielding a corresponding division rate.

Further observations are consistent with this picture. First, 1/*T*_*cyc*_ changes on a timescale (within 25 minutes of the shift) that is shorter than the cell cycle duration, or even the C+D period (∼65-75 minutes) [22]. Scenarios in which division occurs a fixed time after the moment of replication initiation [23–25], with ppGpp affecting the added size by modulating the initiation rate, are inconsistent with these observations. These scenarios require completion of ongoing cell cycles before registering ppGpp effects. The 1/*T*_*cyc*_ response would then exceed the C+D period and decrease exclusively, to match the new *μ*. Second, the growth rate independent model predicts the 1/*T*_*cyc*_ increase is too small to detect in minimal media (Fig. 3D and E), owing to the lower *µ*, which is indeed observed in the experiments (Fig. 3F). Third, upon Mesh1-CFP induction, *T*_*cyc*_ increases within 20 minutes and then stays constant (Fig. S3B). These results indicate that ppGpp can exert control over division, and hence over cell size, in a way that is not mediated hierarchically through its effects on initiation and the rate of growth.

## Discussion

Elucidating the coordination between cell growth and cell cycle progression is a foundational challenge of microbial physiology. The adder mechanism has emerged as a key principle in recent years: it is observed across diverse domains of life and experimental conditions, and elegantly explains how size remains constant despite (division) stochasticity and exponential volume growth [2–4,15,23–27]. By *adding* a constant size every cycle, stochastic size variations are averaged out without needing a specific response to size deviations. How the adder mechanism relates to ppGpp is central, given the inherently intertwined nature of size and growth, yet remains poorly understood. Indeed, ppGpp is implicated in diverse cellular metabolism and growth processes, including regulating ribosome production [28–30], modulating membrane synthesis [31,32] DNA replication initiation [5,33–35], and triggering the stringent response [36–38], a stress reaction that arrests growth until conditions improve.

Here we found that ppGpp is a cell division regulator, and hence serves as a link between growth and size control mechanisms. More specifically, we showed that *E. coli* cells do not follow a hierarchical model, in which cell size adjusts to the growth rate (as in the general growth law [1,17–19], and ppGpp controls growth (by tuning ribosome production depending on amino acid availability, for instance). Rather, the (added) size correlates with the level of ppGpp (instead of the growth rate), and adjusts rapidly to ppGpp deviations, prior to the growth rate response, indicating that ppGpp exerts independent control over cell division and growth.

The findings lead to a number of speculations and implications. ppGpp is known to reflect diverse signals, including nutritional conditions, stress, biosynthetic activity, and metabolite availability [39,40]. The proposed mechanism thus allows division control to integrate a wide spectrum of relevant growth factors. Indeed, these factors may influence division through ppGpp rather than in direct manner, given that division was here found to respond to ppGpp changes alone, under constant nutritional conditions, and before growth itself adjusted. In addition, one may speculate that ppGpp-mediated cell cycle control helps to accommodate different physiological limitations. For instance, a ribosome excess may incur a growth cost while also favoring larger cells, rather than the smaller cell size that would be imposed by the strictly positive size-growth correlation in hierarchical models. Consistently, overexpression of a non-functional protein was shown to yield larger cells [41]. The growth adjustments caused by the small ppGpp changes that are the central focus here were also notably slower than those that characterize the stringent response and large ppGpp changes. The former could be caused by ppGpp inhibition of ribosome production, and the resulting dilution of ribosomes by volume growth.

In recent years, diverse mechanisms have been proposed to explain cell size homeostasis and its dependence on growth [2–4,15,23–27]. Our findings suggest these mechanisms are under the control of ppGpp. For instance, the constant added size is proposed to result from the accumulation of a signaling molecule throughout the cell cycle, which triggers division when a threshold is exceeded [24,42]. One may speculate that ppGpp alters the production of this molecule and its threshold. Owing to the central role of ppGpp in metabolism and its many regulatory targets (hundreds of genes [43] and dozens of proteins [44]), ppGpp could control size in many possible ways. It was found that OpgH can suppress FtsZ ring formation depending on the growth rate [45]. One may consider whether OpgH mediates the ppGpp division effects, though it is not a known ppGpp interactor [46]. Nucleoid occlusion mechanisms have also been proposed to explain variations in added size [2]. Nucleoid volume could be modulated by ppGpp-induced decreases in the overall DNA replication initiation [47–49] or transcription rates [50,51].

In conclusion, our findings establish a link between ppGpp, the central signaling molecule in bacterial growth, and cell size homeostasis, which has implications for the diverse cellular components and processes that are involved in these two pivotal cellular control mechanisms.

## Supporting information

Supplemental Information - Modelling

## Acknowledgements

We thank Rebecca McKinzie, Martijn Wehrens, Nicole Imholz and Marek Noga for experimental assistance and helpful discussions throughout the project and Daan J. Kiviet for sharing his mold for the microfluidic device.

## Author contributions

G.B., S.J.T., and F. B. conceived the experiments, F.B. performed the experiments, F. B., M.C.L. and J.G. performed the modelling, all authors contributed to the writing of the manuscript.

## STAR Methods

### Strains and plasmids

E. coli K-12 strains NCM3722 (CGSC# 12355) and ppGpp^0^ (CF10237) which were transformed with a combination of pRelA*-YFP, pMesh1-CFP or pCFP induction plasmids were used in all the experiments as described in the figures. Plasmid pRelA*-YFP was constructed by replacing mCherry on a pBbS2k-RFP plasmid (kanamycin resistance, Tet promoter, sc101** origin) with a DNA sequence encoding the first 455 amino acids of the native RelA gene. YFP fluorophore mVenus was fused to RelA* via a glycine-serine linker using restriction cloning. Similarly, a codon optimized sequence of Mesh1 or CFP replaced mCherry in the plasmid pBbA5a-RFP (ampicillin resistance, lacUV5 promoter, p15A origin) leading to pMesh1 and pCFP. pMesh1-CFP was built similarly to pRelA*-YFP using restriction cloning to fuse CFP with Mesh1 via a glycine-serine linker.

Chemocompotent NCM3722 cells were transformed with the pRelA*-YFP plasmid and spread on LB Agar plates with 25 ug/ml kanamycin. pMesh1-CFP and pCFP plasmids were transformed and plated on LB Agar plates with 50 ug/ml ampicillin. Plates older than 3 weeks were discarded and fresh transformations were prepared to prevent possible mutants.

Chemocompotent ppGpp^0^ cells were co-transformed with pRelA*-YFP and pMesh1-CFP and plated on LB Agar plates with 50 μg/ml ampicillin, 25 μg/ml kanamycin and 100 μM IPTG to allow growth. Without 100 μM IPTG, leakage from pRelA*-YFP inhibits growth enough to prevent visible colonies next day morning (data not shown). Plates with colonies were only used on the day where the colonies first appear (next morning after transformation) as older plates loose viability rapidly.

### Culture conditions

Cells for bulk and microscopy experiments were grown using defined MOPS medium containing 0.2% (with volume) carbon source [52] with 100 μM MnCl_2_ (minimal media). Rich medium is the same as minimal media except for the supplementation of 0.2% Casamino acids, 400 μg/ml serine and 40 μg/ml tryptophan. 50 μg/ml ampicillin and 25 μg/ml kanamycin were added to the media along with appropriate plasmid bearing strains.

For the microscopy experiments, cells were initially inoculated in 10 mL tubes with 5 ml MOPS rich medium from a single colony in the morning. The tube was placed in a 37°C room on an orbital shaker until the growth becomes visible (OD ≈ 0.1). Cells were then spun down at 4000 G for 5 minutes and re-inoculated in 10 µL top media. 2 µL of the concentrated cells were injected into the microfluidic chip by hand via a p2.5 pipette and appropriate pipette tip.

ppGpp^0^ strain requires different handling due to its inability to respond to stress. For the 96 well plate experiments, a single colony was inoculated in 5ml Glucose rich media with 100 μM IPTG and placed on a shaker for up to 4 hours until OD_600_ reaches 0.4. After that 40 ng/ml Dox is added to prime the culture with high ppGpp production which allows it to handle stress and initiate growth in minimal media. This culture is then diluted in fresh media (MOPS minimal with different carbon sources) without any inducers. Immediately after the dilution, 98 μL of the culture is pipetted into wells of 96 well plates together with 1 μL Dox and 1 μL IPTG (both at 100x final concentration), reaching a final volume of 100 μL. 96 well plates which were prepared as above were placed in a BioTek Synergy HTX plate reader @37°C with constant orbital shaking. OD was measured every 10 minutes. A similar method is applied for the microscopy experiments with the ppGpp^0^ strain. A single colony is grown in rich media with proper antibiotics from the morning and when OD reached 0.1, 40 ng/ml Dox is added to induce ppGpp production, which promotes cell survival during chip loading.

The chip is then placed under the objective in a warm chamber set to 37°C for all the experiments. After cells populate the growth chamber input and output tubes are connected to the chip and appropriate media is pushed through at 500 μl/hr. This corresponds to 250-500x dilution per hour since the chip’s inner volume is between 1-2 μL.

### Microfluidic flow cell

The microfluidic chip’s Epoxy mold which was kindly sent by Daan J. Kiviet from Ackermann Lab is a variant of the mothermachine from Jun lab. Each flow line consists of an input which splits up into 2 arms, in each arm there are a number of extruding growth chambers varying in depth and width (80, 60, 40, 20, 10, 5 µm width, 60, 30, 50, 40 µm depth) and a single output after the 2 arms reconnect into a single line.

Chips were built by first preparing the PDMS mix using the protocol from the reference [15]. Polymer and curing agent (Sylgard 184 elastomer, Dow Corning) were prepared by mixing 7.7 g of polymer with 1mL of curing agent. The slight deviation from the suggested 10 g per 1 mL was implemented to create a more rigid chip allowing low height growth chambers to remain intact. Mixture was then thoroughly mixed using a vortexer and a plastic mixer. Then the mix was poured into the Epoxy mold (provided by Ackermann lab) and placed in a desiccator for 30 minutes to remove air bubbles formed during mixing. Then the mold is baked at 80°C for 1hr. After the baking period, the PDMS chip was removed from the mold using scalpels and rough edges were cut to allow for better binding to glass. Inlet and outlet holes were punched using a hole puncher. The PDMS chip was then covalently attached to a glass slide by using a hand-help corona treatment device (model BD-20ACV, Electro-Technic Products). Application was done by passing the corona treatment device 6-7 times, each pass lasting ∼5 seconds, 5-10 mm away from the surface of both the PDMS chip and glass cover slip. After the corona application, PDMS chips were placed on the treated glass surface and tapped by a gloved finger to assure full bonding. Prepared chips could be used couple weeks after preparation however after more than a month, chambers start to collapse, therefore chips older than 1 month were not often used.

### Imaging and Image Analysis

Cells growing in the microfluidic chambers were imaged using an inverted Nikon TE2000. Using 100x and a 1.5x zoom lenses in tandem a pixel size of 0.041 µm was achieved. Imaging was done using a CMOS camera (Hamamatsu Orca Flash 4.0) via illumination from an LED light source (ImSpec, HPX-L5) with a liquid light guide. The microscope stance was equipped with a computer-controlled stage (Marzhauser, SCAN IM 120 3 100) allowing the stage to move between several chambers for imaging. A phase contrast image was taken every minute and a fluorescence image every 5 minutes using Chroma filter set 49003 and a computer-controlled shutter (Sutter, Lambda 10-3 with SmartShutter). Control of the automated microscopy systems was achieved through MetaMorph software. Each experiment lasts between 24-36 hours.

Images were initially visually checked for issues such as cells washing away from the wells or halting of growth due to clogs. After the initial checks, a MatLab based software customized by the Tans Lab was used to quantify growth rates and cell sizes. Individual cells were identified and tracked from phase contrast images. Cell’s lengths and volumes were estimated assuming the shape of cylinders with semicircular caps and fitting a polynomial to skeletons of binary cell masks. Estimated length data through time was used to calculate instantaneous growth rate and average growth rate using exponential fits. Fluorescence values were calculated from a strip inside the cell area to decrease errors caused by fluorescence falloff that occurs at the edges of the cell. Added length was calculated by subtracting length at division (end of cycle) from length at birth (beginning of cycle). Duration of the cycles were calculated as the time between birth and division. Any cell that did not divide within the growth chamber was ignored along with cells that approached the exits of the wells due to tracking issues caused by increased cell speeds near the exit.

### Calculation of stabilization time

Data points within a sliding window of width ∼*T*_*cyc*_ were compared against the data points where growth rate became stable after induction. As the sliding window moves through time, both growth rate and added length decrease and approach the final stable value. When the t-test between the data points in the sliding window and in the final stable growth regime yielded a p-value that exceeded 0.05 in three consecutive sliding time steps, the center of the window defined the “stabilization time”. For each experiment the data was randomly split into 3 parts and each part was analyzed using the same method (black dots, Figure 2D).

### Division frequency analysis

Box plots (Figure 3D-F) were determined for measured values of 60/*T*_*cyc*_, in a period between 150 min. before induction and the moment of induction (green), and in a period between the moment of induction until two times the average pre-shift cell cycle time (blue).

### Models

We considered two alternative models describing size control during the transient. In both models a cell divides with a hazard rate [21] that depends on the added size and a target added size ΔL (which in stationary conditions corresponds to the average added size). Both the target added size ΔL(t) and the growth rate µ(t) are functions of time during the transient. In both the models the growth rate µ(t) is exponentially relaxing to the stationary value observed in each of the experiments.

The two models differ for the relation between growth rate and size scale during transient. In the hierarchical model, the typical size is a deterministic function of the growth rate (ΔL(t) = D*exp(µ(t)*T). Parameters were as follows: T = log(*ΔL*_*Final*_/*ΔL*_*Initial*_)/(*µ*_*Final* -_ *µ*_*Initial*_) and D = *ΔL*_Initial_*exp(-*µ*_*Initial*_*T). In the direct model, ΔL(t) is relaxing to the stationary value linearly in two pre-shift cycle duration time.

## Supplemental Information

**Supplementary Figure 1:**
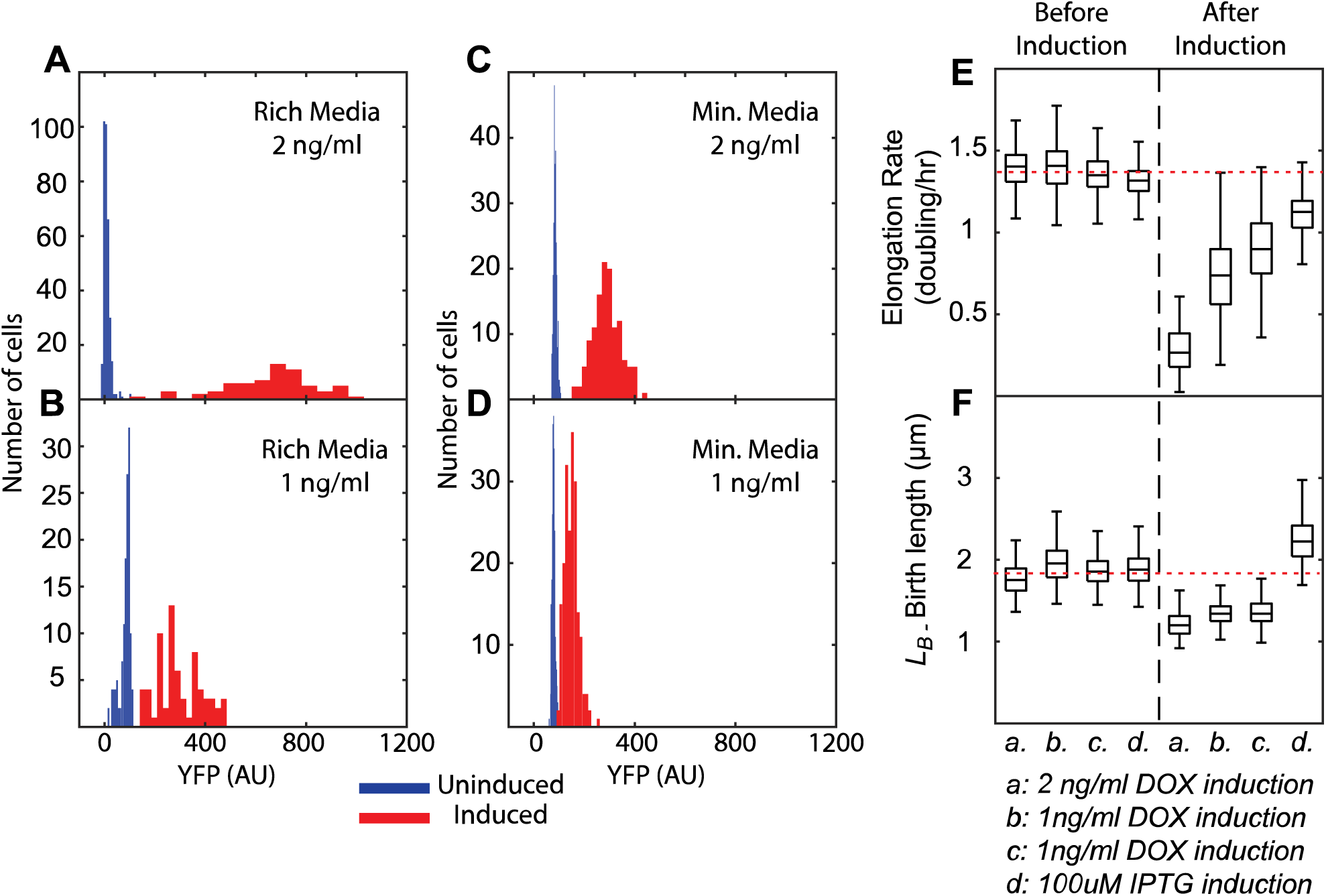
Expression, growth, and size characterization. **(A-D)** Histograms of cellular fluorescence for different conditions. For all indicated induction and growth conditions, fluorescence reporter distributions show that the cellular populations are induced homogeneously. Cells were grown in either rich (with amino acids) or minimal (without amino acids) MOPS-Glucose media. Fluorescence values of individual cells were measured before or after induction of RelA*-YFP with 1 or 2 ng/ml Dox. **(E-F)** Cellular growth rates and birth sizes for different strains and conditions. The strains carry one or two plasmids that encode: RelA*-YFP, inducible by Dox (*a*-*b*), RelA*-YFP, inducible by Dox, and CFP, inducible by IPTG (*c*) or Mesh1-CFP, inducible by IPTG (*d*). The cells were grown in MOPS minimal media in a microfluidic device. Cells were characterized before and after RelA*-YFP or Mesh1-CFP induction, in terms of their growth rates (panel E) and birth lengths (panel F). For strain *c*, CFP was induced always with 200 μM IPTG; both before and after RelA*-YFP induction. The results show minimal growth rate changes caused by CFP expression only (a,b vs c, left). After the induction of RelA*-YFP or Mesh1-CFP, all the strains grow slower, as discussed in the main text. Induction of Mesh1-CFP with 100 μM IPTG leads to an increase of cell size, which is not observed for CFP induction even at 200 μM IPTG (d, right vs c, left, respectively), indicating the size effect is caused by Mesh1-CFP rather than CFP. 2 ng Dox induction decreases the growth rate to ∼0.25 db/hr (a, right). Consistently, 1 ng Dox induction leads to a higher final growth rate that is similar for b and c. Red dotted horizontal line shows the average of 4 experiments before induction of RelA*-YFP or Mesh1-CFP. Number of cell cycles measured: N=257, 264, 355, 403, 150, 316, 389, 241 left to right (panels E and F).

**Supplementary Figure 2:**
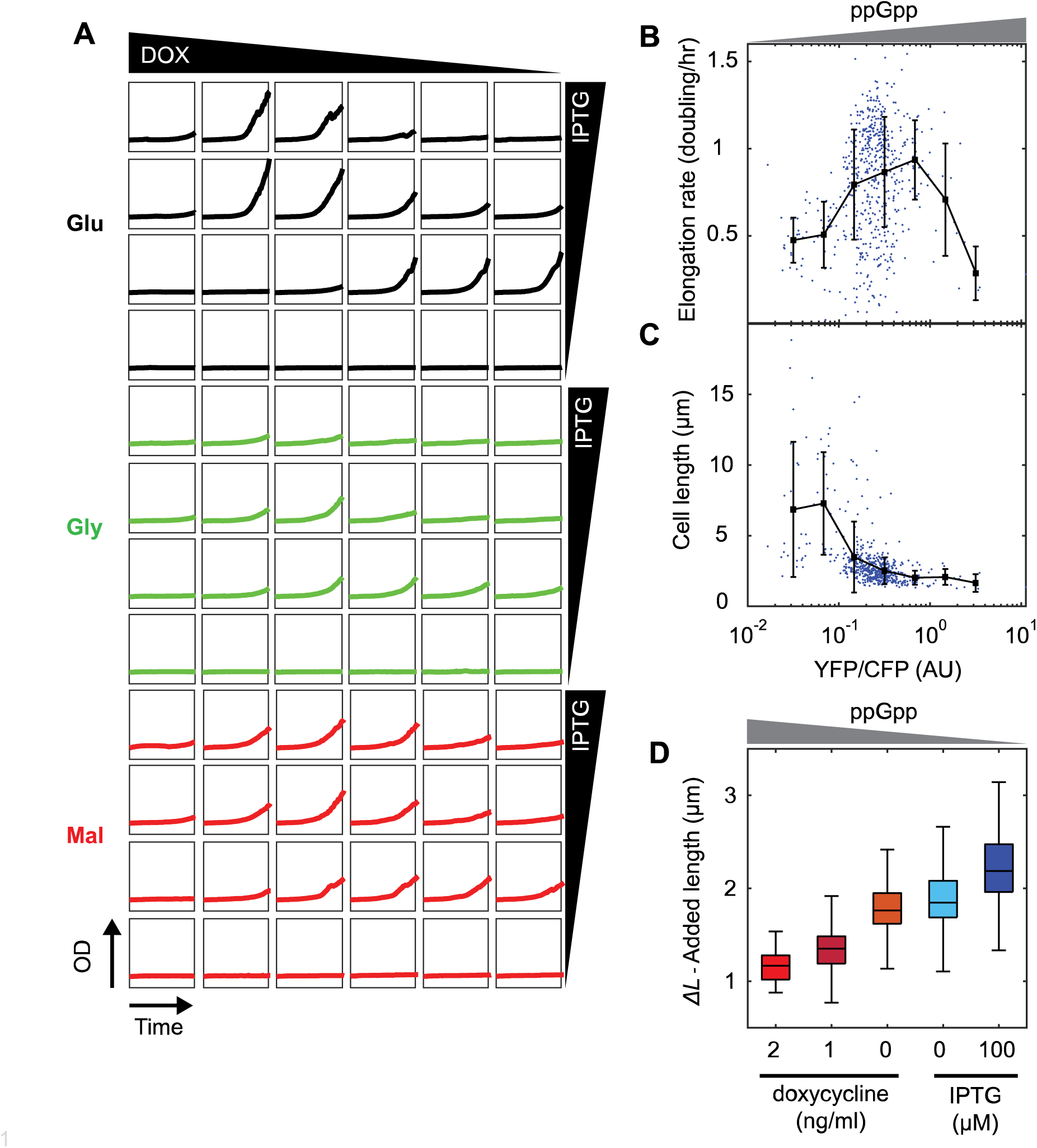
Dependence of cell growth and size on RelA* and Mesh1. **(A)** Optical density (OD) measurements (range 0-0.5 AU) in time (range 0-900 min) for a ppGpp^0^ (ΔRelA, ΔSpoT) strain that is co-transformed with both the pRelA*-YFP (ppGpp synthesis) and pMesh1-CFP (ppGpp hydrolysis) plasmids, grown in minimal media with different carbon sources (black-glucose, green-glycerol, red-malate). Without Mesh1-CFP induction by IPTG (bottom row) no growth is detected. This is consistent with expression from the pRelA*-YFP plasmid being lethal, due to the resulting accumulation of active ppGpp. Similarly, if Mesh1-CFP is maximally induced with IPTG (maximal hydrolysis) while RelA*-YFP is not induced with Dox (minimal synthesis), growth is minimal (top right plots), as the resulting lack of ppGpp is detrimental to growth in minimal media. Growth effects caused by induction of one of the enzymes can be compensated by induction of the other enzyme, as shown by the diagonal boxes (top-left to bottom-right) displaying the fastest growth. This result is consistent with RelA*-YFP synthesizing ppGpp, and Mesh1-CFP hydrolyzing it. **(B)** Growth rates of single cells against ppGpp level, as quantified by YFP/CFP, for a ppGpp^0^ strain with pRelA*-YFP and pMesh1-CFP plasmids. Variation along YFP/CFP axis indicates stochasticity of RelA*-YFP and Mesh1-CFP expression. Growth shows peak for increasing YFP/CFP, while the (birth) size decreases systematically (panel C). These findings support similar findings in main text Fig. 1C-E. **(C)** Birth size of single cells against ppGpp level, as quantified by YFP/CFP. At low ppGpp (low YFP/CFP), cells grow slower yet are larger. **(D)** *ΔL* increases systematically with decreasing ppGpp levels (in similar vein as the birth size in Fig 1D), rather than with the growth rate, which peaks with ppGpp (Fig. 1C). Experimental details: see Figure 1 C-D.

**Supplemental Figure 3.**
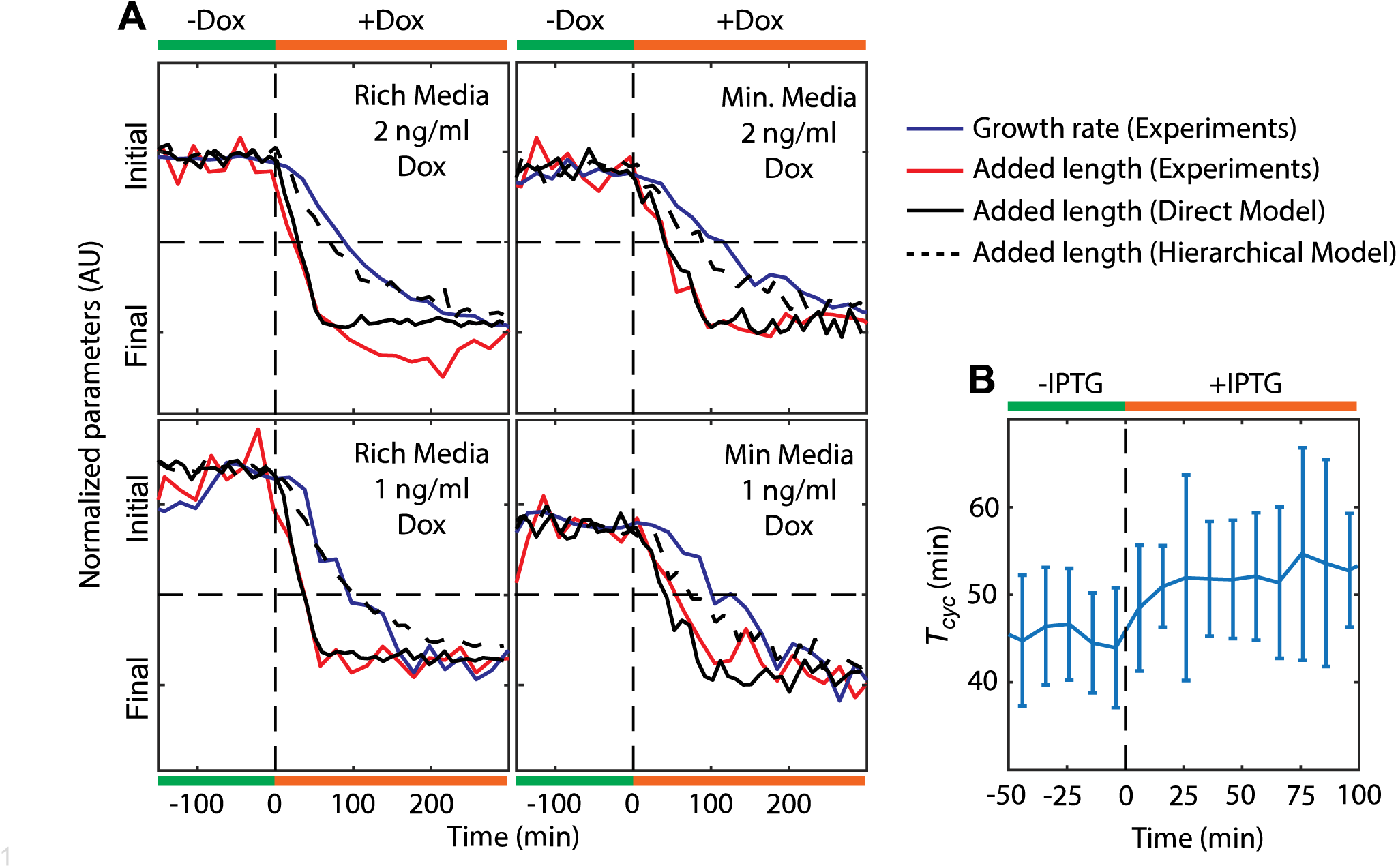
Cell cycle parameters during ppGpp shifts of different size in different media. **(A)** Normalized cellular growth rate and added size during ppGpp shifts. A strain carrying RelA*-YFP was grown in rich (with amino acids) or minimal (without amino acids) MOPS-Glucose media. Induction of RelA*-YFP with 2 or 1 ng/ml Dox occurs at T = 0 mins (vertical dashed line). Indicated are moving averages of the growth rate (blue curve) and added length (red curve) from experiments, and computed added lengths from two models (solid black curve: direct model, dashed black curve: hierarchical model). The objective here is to assess the speed of adjustment from initial to final values. Hence, the data sets were normalized to all start at the same position on the Y-axis, and to also end at the same position (Initial and Final, respectively). Horizontal dashed lines are half-way between initial and final values. Hierarchical adder model does not follow the experimentally observed ΔL trajectory (red curve), while the direct adder model does show a close match. **(B)** Cell cycle duration during ppGpp down-shift. Induction of Mesh1-CFP (T=0 mins, vertical dashed line) leads to rapid increase in *T*_*cyc*_ suggesting inhibition of division by decreased ppGpp levels.

