## Supplemental Information - Modelling for "ppGpp is a bacterial cell size regulator"

### ppGpp is a bacterial cell size regulator
### Supplementary text — Theoretical framework

#### I. HIERARCHICAL AND DIRECT MODELS OF DIVISION CONTROL.

This section describes in mathematical terms the theoretical framework for the hierarchical and direct models of division control used in the Main Text.

##### Notation and model definition.

We assume that in steady conditions cells grow exponentially with (average) elongation rate $\mu$

$$\frac{dL(t)}{dt} = \mu L(t) \ , \tag{1}$$

where $L$ is cell length. It naturally follows that, despite $\mu$ may fluctuate, the average cell length grows as $L(t) = L_B e^{\mu(t-T_B)}$, where $L_B$ is the size at birth and $T_B$ is time at birth.

For a given $\mu$, we model cell division using the hazard-rate framework [1], which describes the probability per unit time that a cell with size $L$, initial size $L_B$, and elongation rate $\mu$ divides. In general, one can write the probability that a cell born at time $T_B$ with size $L_B$ divides at time $T_B + T_{cyc}$ as

$$p(T_{cyc}|L_B, \Delta L, \mu, T_B) = -\frac{d}{dT_{cyc}} \exp\left(\int_{T_B}^{T_B+T_{cyc}} dt \ \mu \ g_\sigma(L(t)|L_B, \Delta L, \mu, t)\right) \ , \tag{2}$$

where $g_\sigma$ is related to the hazard rate [2]. Under steady conditions, $g_\sigma$ does not depend on $t$. Moreover, the collapse of the size distributions across conditions imply that the dependence of $g_\sigma$ on $\mu$ [2] can be reabsorbed into the dependence of $\Delta L$ on $\mu$, and Eq. 2 reads

$$p(T_{cyc}|L_B, \Delta L, \mu, T_B) = -\frac{d}{dT_{cyc}} \exp\left(\int_{T_B}^{T_B+T_{cyc}} dt \ \mu \ g_\sigma(L(t)|L_B, \Delta L(\mu))\right) \ , \tag{3}$$

where the function $\Delta L(\mu)$ encodes the dependency of the typical size on the elongation rate, known as Schaecter's law [2–4].

Following [2] we use

$$g_\sigma(L|L_B, \Delta L) = \frac{1}{\sqrt{2\pi}\sigma} \frac{\exp\left(-s^2\right)}{1 - \mathrm{Erf}(s)}\Bigg|_{s=\frac{\log(L)-\log(L_B+\Delta L)}{\sqrt{2}\sigma}} \ . \tag{4}$$

While this definition might appear cumbersome, it corresponds to an adder model with lognormal noise, which well describes single-cell data in steady growth conditions [2, 4]. The parameter $\Delta L(\mu)$ is the average added size and $\sigma$ is the standard deviation of the log-added size.

###### *Generalization to non-steady growth conditions*

Under non-steady growth conditions — in our case, when the elongation rate is changing in response to changing ppGpp levels — $\mu(t)$ is itself time dependent (in average and fluctuations). The average growth curve of a cell is therefore the solution of

$$\frac{dL(t)}{dt} = \mu(t) L(t) \ , \tag{5}$$

which reads

$$L(t) = L_B \exp\left(\int_{T_B}^{t} ds \ \mu(s)\right) \ , \tag{6}$$

for $t > T_B$. Under non-steady growth, the model has to take explicitly into account the dependency of the hazard rate, appearing in Eq. 2, and of the other parameters on $T_B$.

*Hierarchical model*

Under the hierarchical model, the mechanisms affecting the added size are downstream of the ones affecting the elongation rate. In this case equation 3 reads

$$p(T_{cyc}|L_B, \Delta L, \mu, T_B) = -\frac{d}{dT_{cyc}} \exp\left(\int_{T_B}^{T_B+T_{cyc}} dt\ \mu(t)\ g_\sigma\left(L(t)|L_B, \Delta L(\mu(t))\right)\right)\ , \tag{7}$$

where $T_B$ is the time at birth of a given cell and $L(t)$ is given by equation 6.

*Direct model*

Contrarily to the hierarchical model, in the direct model the dynamics of the elongation rate $\mu(t)$ does not constrain the time dependency of $\Delta L$. The probability of division is therefore

$$p(T_{cyc}|L_B, \Delta L, \mu, T_B) = -\frac{d}{dT_{cyc}} \exp\left(\int_{T_B}^{T_B+T_{cyc}} dt\ \mu(t)\ g_\sigma\left(L(t)|L_B, \Delta L(t)\right)\right)\ , \tag{8}$$

where $\Delta L(t)$ encodes the dependency of the added size on time.

**The direct model predicts initial increase of $1/T_{cyc}$**

This section presents an analytical argument applied to the direct model showing the mechanism that gives rise to an initial increase of $1/T_{cyc}$ described in Fig. 3 of the main text.

Before the shift, at time $t < 0$, an average cell adds a constant size $\Delta L(0)$, is born with size $L_B(0) = \Delta L(0)$, grows with elongation rate $\mu(0)$, and divides after a time $T_{cyc}(0) = \log 2/\mu(0)$ from birth. In steady growth, size and elongation rate are related via Schaecter's law, which we approximate as

$$\Delta L(0) = \Delta^* \exp\left(\mu(0)T^*\right)\ , \tag{9}$$

where $\Delta^*$ and $T^*$ are estimated as described in the main text. If $\Delta L(\infty)$ and $\mu(\infty)$ are the added size and growth rate after the shift, we have that

$$\Delta L(\infty) = \Delta L(0) \exp\left((\mu(\infty) - \mu(0))T^*\right)\ . \tag{10}$$

For sufficiently large times after the shift, the added size and the elongation rate relax to the final values $\Delta L(\infty)$ and $\mu(\infty)$. The division time before (after) the shift is given by $T_{cyc}(0) = \log(2)/\mu(0)$ ($T_{cyc}(\infty) = \log(2)/\mu(\infty)$). In our case $\mu(\infty) < \mu(0)$ and therefore $T_{cyc}(0)/T_{cyc}(\infty) = \mu(\infty)/\mu(0) < 1$. Figure 3 shows that in the experiments, at the beginning of the shift, the values of the division rate are growing. If we define $1/T_{cyc}(t)$ to be the division rate during the shift, we observe that $T_{cyc}(0)/T_{cyc}(t^*) > 1$, approximately for $t^* < 2T_{cyc}(0)$ (i.e., within the first two cell cycles after the shift).

According to the direct model, the dynamics of the added size and the elongation rate are decoupled during the shift. In particular the added size $\Delta L$ relax change more rapidly than the elongation rate $\mu$. For every cell, one has the constraint that

$$L_B + \Delta L = L_B \exp\left(\mu T_{cyc}\right)\ ,$$

hence

$$\log\left(1 + \frac{\Delta L}{L_B}\right) = \mu T_{cyc}\ . \tag{11}$$

At steady state, on average $\Delta L(0)/L_B(0) = 1$, but this number decreases immediately after the shift due to the faster decrease of $\Delta L$. We can assume that there is a moment just after the shift where cells still grow approximately with elongation rate $\mu(0)$, but their added size to initial size ratio has already decreased, hence

$$\mu(0)T_{cyc}(t^*) = \log\left(1 + C\right)\ , \tag{12}$$

with $C < 1$. By using $\mu(0)T_{cyc}(0) = \log 2$, we obtain

$$\frac{T_{cyc}(0)}{T_{cyc}(t^*)} = \frac{\log 2}{\log(1+C)} > 1 \ . \tag{13}$$

In other words, $T_{cyc}(t^*) < T_{cyc}(0)$, i.e., the division rate has to initially increase with respect to its old steady-state value, before it can decrease to its new steady-state value in the high ppGpp growth condition.

**The rapid decrease of $\Delta L$ and the initial increase of $1/T_{cyc}$ is incompatible with initiation determining division.**

This section presents a theoretical argument against the hypothesis that ppGpp acts on cell division through replication initiation. Specifically, we will argue that a model where the change in added size across the shift is modulated by initiation rate through a replication-limited cell-division process is incompatible with the experimental data. Assuming this replication-limited model, before the shift replication initiation occurs at a typical size $L_I(0)$ and the corresponding division follows after a time $T_{C+D}$, of the order of $65 - 75$ minutes. In steady growth conditions,

$$2^n L_B(t + T_{C+D}) = L_I(t)\exp(\mu T_{C+D}) \ , \tag{14}$$

where $n$ is the number of overlapping rounds of replication (in slow growth $n = 1$) and where $t$ stands for absolute time. On average, the initiation of replication in a cell at time $t$ determines division at time equal to $t + T_{C+D}$. It easy to observe that

$$\Delta L(t + T_{C+D}) = \Delta I(t)\exp(\mu T_{C+D} - n\log 2) \ , \tag{15}$$

where $\Delta I$ is the added size between two consecutive initiations. In steady growth, the average $\Delta L$ and $\Delta I$ are time independent and Eq. 15 sets their relationship.

Following a shift, the initiation rate and the elongation rate change, while replication speed and the time $T_{C+D}$ can be assumed constant (or at least bounded from below). This implies that, if division was set by initiation, any change in the added size $\Delta L$ would be observable after a time $T_{C+D}$ from the change of initiation rate. In the experiments, the added size change on a timescale of the order ot 25 minutes, which is incompatible with the scenario of initiation determining division and $T_{C+D} > 60$ minutes.

#### II. CODE TO SIMULATE THE MODELS

Here we report the general structure of the code (written for Python) for both the models.

---

```
import scipy as sp
import numpy as np


def mu_time( t, t0 = 0, tau = transition_timescale_grate ):
    if t < t0:
        return mu1
    elif t >= t0:
        return mu2 + (mu1-mu2) * np.exp(-t/tau)

def size_time( t, t0 = 0, tau = transition_timescale_size ): # not used in the hierarchical model
    if t < t0:
        return size1
    elif t >= t0 and t < t0+tau:
        return size1 + (size2-size1) * (t-t0)/(tau-t0)
    else:
        return size2

def dmu( t, mu , cvmu, dt ):
    mumean = mu_time( t )
    dadt = -( mu - mumean )/timecorrmu + mumean * cvmu * np.random.normal() / np.sqrt(dt * timecorrmu)
    return dadt
```

```python
def hazard_rate( x, x_mean, sigma ):
    q = np.log(x)
    q_m = np.log(x_mean)
    s = (q-q_m) / sigma / np.sqrt(2.)
    h = np.exp( -s*s ) / math.erfc(s)
    h = h * np.sqrt(2./np.pi) / sigma / x
    return h


def check_if_dividing( x, x_mean, sigma, dx ):
    if sigma > 0:
        h = hazard_rate( x, x_mean, sigma )
    else:
        h = 1.*( x > x_mean ) / dx
    if( h*dx > np.random.rand() ): return True
    else: return False




# mu1 pre shift, mu2 post shift
Tcyc1 = 27.1 # minutes
Tcyc2 = 120.6
mu1 = np.log(2.)/Tcyc1 # mu is growth rate base e. #np.log(2.)/Tcyc, Tcyc is the average division time
mu2 = np.log(2.)/Tcyc2
# size1 pre shift, size2 post shift
size1 = 2.19 # micron
size2 = 1.25
## get parameters for schaecter law
schaecter_slope = np.log( size2 / size1 ) / (mu2-mu1)
schaecter_intercept = size1 * np.exp(-mu1 * schaecter_slope )

# transition timescales
transition_timescale_grate = 111.6 # [min] # characterisitc timescale of transition in growth rate
transition_timescale_size = 2*Tcyc1 # characterisitc timescale of transition in added size # not used in
    the hierarchical model

timecorrmu = 10 #[min] # autocorrelation time of growth rate dynamics

t_initial = -1000. # initiial time of simulation in min
t_final = 750. # final time
sigma = 0.1 # coefficient of variation of size
cvmu = 0.074 # coefficient of variation of single-cell growth rate

fout = open( "outputfile.dat" , "w" )
fout.write( "cell t t0 xf x0 gratemean\n" )

dt = 0.1 # [min]
Ncells = 1000
for i in range(Ncells):
    print(i)
    t = t_initial + (np.random.rand()-0.5)*100 # desync initial cell population
    mu = mu_time( t ) * (1. + cvmu * np.random.normal() )
    x0 = size_time(t)
    x = x0
    t0 = t
    LastDivision = True
    while t < t_final or LastDivision:
        mumean += mu
        npoint += 1
        t = t + dt
        dx = mu * x * dt
        mu = mu + dmu( t, mu , cvmu, dt )*dt
```

```
    x = x + dx
    x_mean = division_size_control( x0, mu, typecontrol, t ) # this function differs between the two
        models
    if( check_if_dividing( x , x_mean, sigma, dx ) ):
        fout.write( "%d %f %f %f %f %f\n" % (i, t, t0, x, x0, mumean/npoint) )
        t0 = t
        x0 = x * 0.5
        x = x0
        if ( t > t_final ): LastDivision = False

fout.close()
```

**Hierachical model**

```
def division_size_control( x0, mu, typecontrol, t ):
    delta_x = schaecter_intercept * np.exp( mu * schaecter_slope ) # this choice defines the hierarchical
        model
    xf = x0 + delta_x
    return xf
```

**Direct model**

```
def division_size_control( x0, mu, typecontrol, t ):
    delta_x = size_time( t ) # this choice defines the direct model
    xf = x0 + delta_x
    return xf
```


[1] Matteo Osella, Eileen Nugent, and Marco Cosentino Lagomarsino. Concerted control of Escherichia coli cell division. *Proceedings of the National Academy of Sciences of the United States of America*, 111(9):3431–5, mar 2014.

[2] Jacopo Grilli, Matteo Osella, Andrew S. Kennard, and Marco Cosentino Lagomarsino. Relevant parameters in models of cell division control. *Physical Review E*, 95(3):032411, mar 2017.

[3] Hai Zheng, Yang Bai, Meiling Jiang, Taku A. Tokuyasu, Xiongliang Huang, Fajun Zhong, Yuqian Wu, Xiongfei Fu, Nancy Kleckner, Terence Hwa, and Chenli Liu. General quantitative relations linking cell growth and the cell cycle in escherichia coli. *Nature microbiology*, May 2020.

[4] Suckjoon Jun and Sattar Taheri-Araghi. Cell-size maintenance: universal strategy revealed. *Trends Microbiol*, 23(1):4–6, Jan 2015.
